# Strategies for improved xylitol production in batch fermentation of sugarcane hydrolysate using *Saccharomyces cerevisiae*

**DOI:** 10.1101/2022.05.25.493426

**Authors:** Frank Uriel Suarez Lizarazo, Gonçalo Amarante Guimarães Pereira, Fellipe da Silveira Bezerra de Mello

## Abstract

A plethora of studies have focused on improvements of xylitol production. The challenges of establishing a biotechnological route for the industrial production of this sugar have been explored using different microorganisms and renewable feedstock. Nevertheless, sugarcane biomass has been neglected as the pentose source for xylitol production using *Saccharomyces cerevisiae*. Therefore, here we investigate the use of an industrial *S. cerevisiae* strain for xylitol production in batch fermentation of non-detoxified sugarcane straw hydrolysate, envisioning the diversification of the current infrastructure used for second-generation bioethanol production from the same lignocellulosic material. In order to optimize the xylose conversion in a non-fed cultivation system, guidelines in cell inoculum and medium supplementation are suggested, as well as the first attempt to use electro-fermentation for this purpose. Accordingly, our results show that the increase in initial cell density and hydrolysate supplementation allows a xylitol production of 19.24 ± 0.68 g/L, representing 0,132 g/L.h productivity.

## 1. INTRODUCTION

Xylitol is a naturally-occurring biomolecule present in certain fruits and vegetables, and also as a byproduct of carbohydrate metabolism in humans (Rehman et al., 2016). This five-carbon sugar alcohol has a sweetness equivalent to sucrose while presenting only 60% of its calorie content, making it an appealing natural sweetener. Besides the well-established application in the food and beverage industry (Dasgupta et al., 2017), xylitol also finds other commercial applications due to its vast beneficial properties, including: low glycemic index (Albuquerque et al., 2014), noninvolvement in the insulin metabolic pathway (Rafiqul and Sakinah, 2013), high cooling power (Dasgupta et al., 2017), and anti-cariogenic effect (Ur-Rehman et al., 2015). With a worldwide market value in 2021 of USD 471.19 million (Grand View Research, 2021), the xylitol industry represents a hotspot for investments and its revenue is expected to grow in the next years.

Large-scale commercial production of xylitol is currently performed through the hydrogenation and reduction of xylose, purified from xylan present in hemicellulosic hydrolysates. The industrial process encompasses a harsh chemical operation, in which the feedstock undergoes high pressure and temperature conditions in a catalytic-driven reaction, figuring as an energy-intensive industry (Felipe Hernández-Pérez et al., 2019). Xylitol production improvements are paramount for the commercial expansion of such added-value molecule (Albuquerque et al., 2014), especially in a world with raising environmental awareness. In this matter, prospects for biotechnological alternatives have been extensively investigated, since it avoids the catalyst financial and environmental burden, circumvents the need for substrate purification, and requires milder conditions (Queiroz et al., 2022).

A promising option for the bioproduction of xylitol is given by articulating the use of microorganisms, such as yeasts, that may or may not present endogenous xylose metabolism pathways. Studies for xylitol production have been reported with the use of the *Candida* genus and modified *Saccharomyces cerevisiae -* in which genes encoding enzymes involved in pentose consumption have been inserted (Granström et al., 2007). In *S. cerevisiae*, the overexpression of endogenous *GRE3*, or heterologous expression of a xylose reductase, such as *XYL1*, are the main strategies to endow this yeast with improved xylitol production. Although *Candida* spp. shows great natural xylitol productivity, it’s considered an opportunistic pathogen, limiting its use in commercial applications (Granström et al., 2007). Meanwhile, *S. cerevisiae* is a generally recognized as safe (GRAS) microorganism and its well-studied genome makes it the ideal host for genetic engineering (Guirimand et al., 2016). The last also exhibits advantageous traits for industrial fermentation: resistance to low pH, high temperatures, shear stress, and tolerance to aldehyde inhibitors, such as 5-hydroxymethylfurfural (HMF) and furfural (Kogje and Ghosalkar, 2016; Parapouli et al., 2020).

Xylitol production in crude hemicellulosic hydrolysates using *S. cerevisiae* has been reported using birch wood, corn, and wheat as natural xylose sources (Delgado Arcaño et al., 2020; Hou-Rui et al., 2012; Jofre et al., 2022). At the same time, *Candida* spp. has been used as a biocatalyst for other neglected lignocellulosic residues from agro-industrial processes, such as sugarcane biomass (Queiroz et al., 2022). Sugarcane straw represents an attractive pentose source, due to its high availability from the sugar and alcohol industry, where it usually bioaccumulates in the field (De Paiva et al., 2009; Rattanapoltee and Kaewkannetra, 2014). Nevertheless, there are scarcely any reports on this inquiry using *S. cerevisiae*, suggesting a bottleneck for scientific and technological advances. A previous trial for xylitol production in batch fermentation of sugarcane straw hydrolysate using recombinant *S. cerevisiae* resulted in low productivity (de Mello et al., 2021) - possibly due to the recalcitrant fermentation conditions.

In this work, we investigate different cultivation conditions for improved xylitol production from sugarcane straw hydrolysate using an engineered *S. cerevisiae* and batch fermentation - operational condition frequently applied in the Brazilian sugarcane second-generation bioethanol (E2G) industry (Dos Santos et al., 2016). Besides being one of the largest crops worldwide, the sugarcane E2G industry is rapidly evolving (Center for Strategic Studies and Management (CGEE), 2017; Dionísio et al., 2021; Lopes et al., 2016), creating opportunities for the manufacturing of other added-value molecules. While prospects for better xylitol productivity have applied fed-batch cultivation (Baptista et al., 2018; Kogje and Ghosalkar, 2017), the use of a non-fed system accounts for a great share of the fermentation system used in the biomass valorization industry, validating such endeavor. Likewise, co-production of xylitol in the E2G industry has been considered as an alternative to improve the profitability of the process (Ravella et al., 2012).

Accordingly, guidelines here suggested for high cell density initial inoculum and medium supplementation allowed better xylitol productivity, while an improvement was also evidenced using electro-fermentation. With this, we present an attractive alternative to face the great demand for energy and carbon footprint generated by the current chemical route for xylitol production, while using the same process infrastructure already available for the E2G industry.

## 2. MATERIAL AND METHODS

### 2.1. YEAST STRAINS

The *S. cerevisiae* strains used in this study are FMYX and CENPKX (de Mello et al., 2021), derived from industrial Brazilian bioethanol strain SA-1 (Basso et al., 2008) and from the laboratory CEN.PK-122 (van Dijken et al., 2000), respectively. Both modified yeasts express the *xyl1* gene encoding a xylose reductase (XR).

### 2.2. SUGARCANE STRAW HYDROLYSATE PRODUCTION

The biomass hydrolysate was prepared using steam-exploded sugarcane straw (23.9% solids), donated by GranBio SA (Bioflex plant), and Cellic CTec3 enzyme (6% w/w glucan) for the enzymatic hydrolysis. The enzyme of choice is a cellulase and hemicellulase complex with efficient enzymatic activity, largely applied in commercial processes (“Cellic® CTec3 HS | Higher biomass conversion yields | Novozymes,” n.d.; Chandel et al., 2019). For each 100 mL of the hydrolysate, 422 μL of the enzyme was used, and pH was adjusted to 5.0 using ammonium hydroxide. Hydrolysis occurred at 250 rpm and 50 °C, for 72 h. The composition of the sugarcane straw hydrolysate used for xylitol production is provided in **Table 1**.

**Table 1.** Sugarcane straw hydrolysate composition used for xylitol production.

| Component | Concentration (g/L) |
| --- | --- |
| Glucose | 42.75 |
| Xylose | 24.39 |
| Glycerol | 1.12 |
| Acetic Acid | 4.97 |
| Furfural | 0.98 |
| 5-hydroxymethylfurfural | 0.65 |

### 2.3. XYLITOL PRODUCTION IN BATCH FERMENTATION

Fermentation of the non-detoxified sugarcane hydrolysate was performed in 150 mL Erlenmeyer flasks with a working volume of 50 mL at 30 °C, 200 rpm, and aerobiosis, for approximately 145 hours. For inoculum, yeast cells were grown overnight in YPD medium (10 g/L yeast extract, 20 g/L peptone, 20 g/L glucose) at 30 ºC and 250 rpm. Initial cell density (OD_600_) of either 1.0 or 2.5 was used - accordingly discriminated throughout the results section. Cultivation also occurred with hydrolysate supplementation with YP (20 g/L of peptone and 10 g/L of yeast extract). Fermentations were conducted in biological triplicates. Samples were withdrawn for HPLC analysis.

### 2.4. XYLITOL PRODUCTION IN AN ELECTRO-FERMENTATION SYSTEM

An additional condition was tested for the industrial strain FMYX, making use of electro-fermentation with a current of 1 Volt. The bioelectric system generates an electric current using a bioelectric cell, and it was constructed based on patent N° WO 2017/108957 Al (International Publication Number). The electro-fermentation was carried out in 600 mL beakers, with a working volume of 300 mL – volume necessary to cover the bioelectric cell. An image of the cultivation system used is presented in the **Figure 2A**. The fermentation system was sealed with plastic wrap. All fermentations were conducted in biological triplicates. Samples were withdrawn for HPLC analysis.

### 2.5. FERMENTATION PARAMETERS CALCULATION AND STATISTICS

Fermentation parameters were defined as follows: xylitol yield (%) as the amount of xylitol produced (g/L) per consumed xylose (g/L); xylitol productivity (g/L.h) as the amount of xylitol produced (g/L) divided by fermentation time (h). All data are presented as mean ± standard deviation. Data treatment was carried out in Excel 2016 Office, where p-value was calculated according to Student’s t-test.

### 2.6. ANALYTICAL METHODS

Glucose, xylose, xylitol, ethanol, furfural, and hydroxymethylfurfural (HMF) concentrations were determined by high-performance liquid chromatography (HPLC), Alliance® HPLC System Waters detectors: refractive index (RI) detector (Water 2414) at 410nm and photodiode array detector (Waters 2998) at 280nm. Column: ion exclusion Aminex HPX-87H column (300 mm × 7.8 mm, BioRad®), temperature, 35°C; eluent, 5 mM sulfuric acid. Equipment from the Genomics and bioEnergy Laboratory (LGE) of the Institute of Biology, Campinas State University (UNICAMP).

## 3. RESULTS AND DISCUSSION

Here, strains CENPKX and FMYX, derived from laboratory CEN.PK-122 (van Dijken et al., 2000) and bioethanol industrial SA-1 (Basso et al., 2008), respectively, were evaluated for their ability to produce xylitol from xylose present in sugarcane straw hydrolysate. The hydrolysate composition used in this work is disclosed in **Table 1**. In order to improve xylitol productivity reported by de Mello et al (2021) – the only publication available where *S. cerevisiae* was used for xylitol production using sugarcane hydrolysate –, different batch fermentation conditions were tested: increased cell concentration for inoculum and hydrolysate supplementation. Finally, the best cultivation guidelines were applied in an electro-fermentation system to evaluate the effect of an electric current in xylitol productivity. **Figures 1** and **2**, and data in **Table 2** compile the results for each of the proposed conditions, displaying concentrations (g/L) of the metabolites consumed or formed during fermentation, and correlated parameters, respectively. For comparison purposes, results previously obtained by de Mello et al (2021) are displayed as the control experiment, where the same strains and hydrolysate were used.

**Figure 1:**
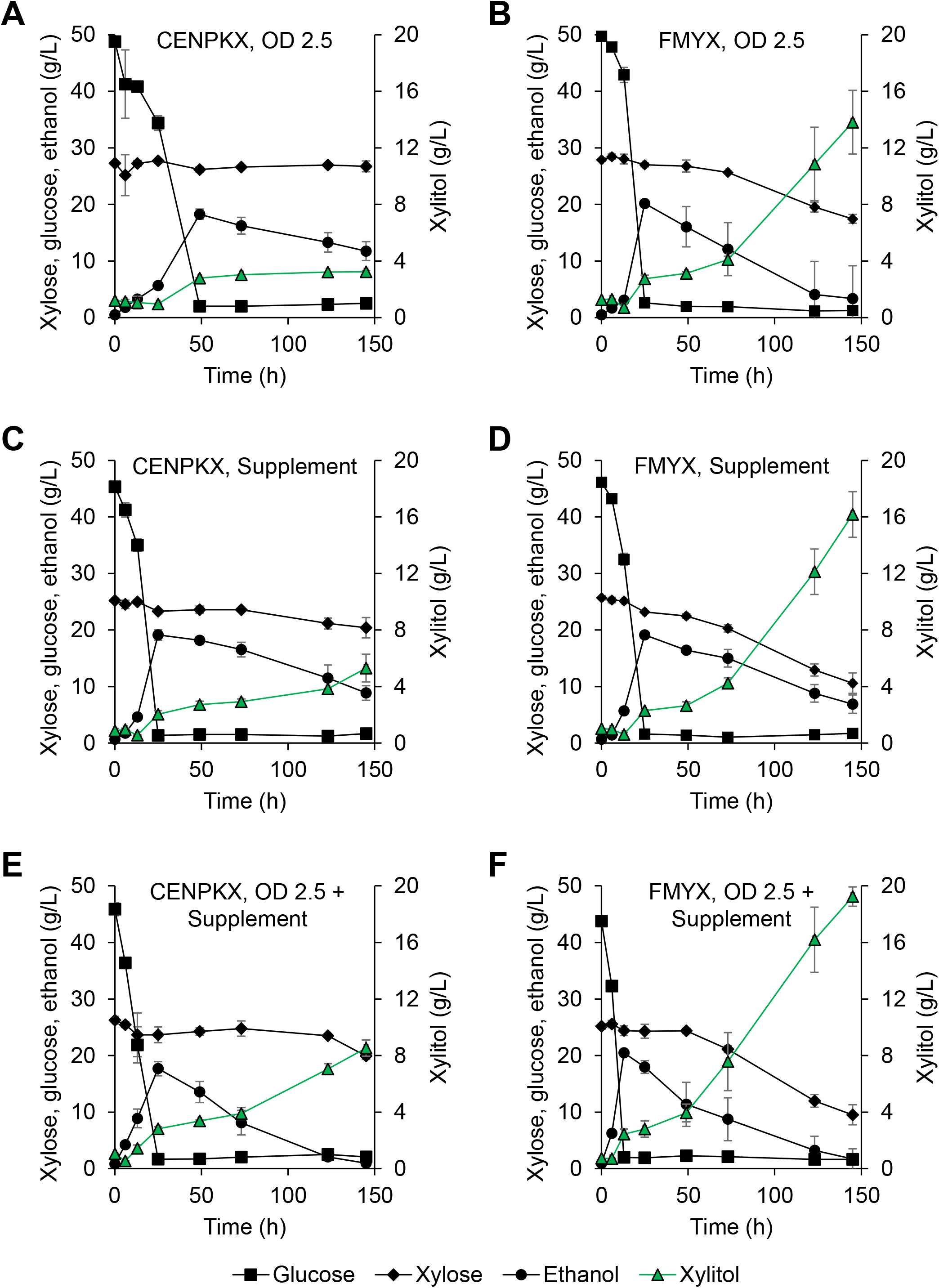
Xylitol production of strains CENPKX and FMYX using sugarcane straw hydrolysate in batch fermentation. Glucose (■), xylose (♦), xylitol (▲) and ethanol (●) curves are represented. Fermentation using high cell density (OD_600_ 2.5) by CENPKX strain (**A**) and by FMYX strain (**B**). Fermentation using supplementation of peptone and yeast extract, by CENPKX strain (**C**) and by FMYX strain (**D**). Fermentation using both conditions (high initial cell density and supplementation of the medium with peptone and yeast extract), by CENPKX strain (**E**) and by FMYX strain (**F**). Errors were plotted as standard deviations and markers represent the average of triplicates for every metabolite.

**Figure 2:**
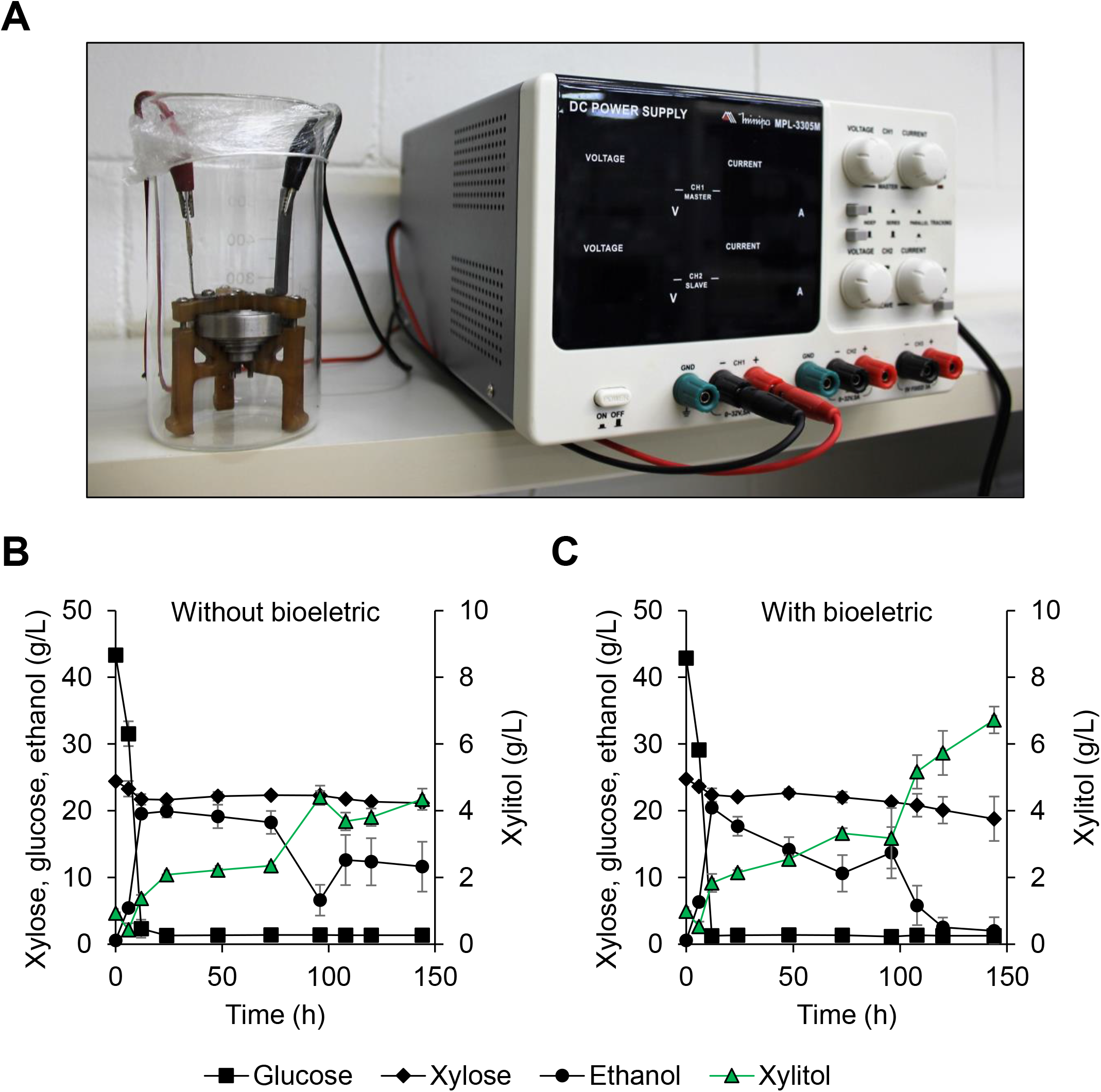
Electro-fermentation of sugarcane straw hydrolysate with high initial cell concentration and media supplementation using strain FMYX. Picture of the system used for the electro-fermentation (**A**). Glucose (■), xylose (♦), xylitol (▲) and ethanol (●) curves are represented in the graphs for conventional batch fermentation in a Becker, as control (**B**), and with the bioelectric cell (**C**). Errors were plotted as standard deviations and markers represent the average of triplicates for every metabolite.

**Table 2.** Xylitol productivity of strains FMYX and CENPKX using raw sugarcane bagasse hydrolysate

| Condition | Titer (g/L) | Yield (g/g) | Productivity (g/L.h) | Reference |
| --- | --- | --- | --- | --- |
|  | FMYX / CENPKX | FMYX / CENPKX | FMYX / CENPKX |  |
| Control | 3,65±0,16** / 1,56±0,45 | 1 / 1 | 0,04 / 0,02 | (de Mello et al., 2021) |
| High cell density | 13,81±2,25** / 3,23±0,1 | 1 / 1 | 0,095 / 0,022 | This study |
| Supplemented | 16,17±1,61*** / 5,30±0,97 | 1 / 1 | 0,111 / 0,036 | This study |
| High cell density + Supplemented | 19,24±0,68*** / 8,52±0,56 | 1 / 1 | 0,132 / 0,058 | This study |
| High cell density + Supplemented + Bioelectric cell | 6,72±0,40** / 4,34±0,31 | 1 / 1 | 0,046 / 0,030 | This study |

\*p-value < 0.05, \*\*p-value < 0.01, \*\*\*p-value < 0.001

Envisioning better xylitol productivity in sugarcane straw hydrolysate, we tested fermentation with higher initial cell density. An initial OD_600_ of 2.5 (equivalent to approximately 8 g/L of cell dry weight) was chosen because this value was reported as an optimal cell concentration for xylitol productivity before it becomes deleterious to xylose metabolism, possibly due to limited aerated conditions created at higher cell densities (Kogje and Ghosalkar, 2017). In this condition, after 145 h fermentation, FMYX was able to produce 13.81 ± 2.25 g/L xylitol, while CENPKX limited its production to 3.23 ± 0,10 g/L of the reduced sugar (**Figure 1A and 1B**). This accounts for 17.45 ± 0.77 g/L xylose left in the medium for FMYX, and 26.73 ± 0.94 g/L for CENPKX after fermentation stopped. In this scenario, 0.095 g/L.h productivity was obtained for the industrial strain, with a 100% yield of xylose conversion. Likewise, other research have reported that greater initial cell density is related to improvement in xylose consumption, suggesting the importance of an adequate inoculum concentration for the fermentation process (Kogje and Ghosalkar, 2017; Matsushika and Sawayama, 2010; Zhong et al., 2009). This phenomenon is possibly explained by the number of cells available to metabolize this pentose (Matsushika and Sawayama, 2010).

Following, we tested whether supplementing the hydrolysate with peptone and yeast extract could improve xylose consumption. For this cultivation status, we preserved a lower initial cell inoculum of OD_600_ 1.0 and added the supplements to the straw hydrolysate. At the end of 145 h fermentation, 16.17 ± 1.61 g/L of xylitol was obtained by the FMYX strain and 5.30 ± 0.97 g/L by the CENPKX strain (**Figure 1C and 1D**). In this scenario, we observed a remaining 10.59 ± 1.81 g/L xylose for the industrial strain and 20.41 ± 1.79 g/L for the laboratory. Concerning productivity, FMYX obtained 0.111 g/L.h and achieved a yield of 100% xylose consumed. These results show that medium supplementation alone allows greater xylitol productivity when compared to higher initial cell density, revealing that this is a key condition for xylitol production in sugarcane hydrolysate.

Supplementation with yeast extract possibly influenced xylose conversion by providing nitrogen source. A similar effect has also been reported in other studies, in which an enhanced xylose conversion was obtained when supplementing the media with yeast extract and peptone, or when combining with solutions containing these nutrients (Bae et al., 2004; Kogje and Ghosalkar, 2017; Preziosi-Belloy et al., 2000; Van Zyl et al., 1989). It is important to note, however, that medium supplementation can generate more efforts in the final purification stages (Parajó et al., 1998).

To evaluate the synergistic effect of both conditions on xylose conversion, we performed a fermentation with both initial OD_600_ of 2.5 and hydrolysate supplementation to test xylitol productivity. At the end of this process, FMYX was able to produce 19.24 ± 0.68 g/L of xylitol, while CENPKX obtained 8.52 ± 0.56 g/L of the reduced sugar (**Figure 1E and 1F**), representing unconsumed xylose of 9.52 ± 1.76 g/L and 19.94 ± 0.63 g/L, respectively. This result represents a productivity of 0.132 g/L.h for the industrial strain, with a xylose conversion yield of 100%. The outcome of this experiment is significantly better than the previously reported 3,65 ± 0,16 g/L of xylitol produced in the same conditions (de Mello et al., 2021). Also, it is noteworthy that the industrial strain outperformed the laboratory in every fermentation condition tested, confirming its robustness towards fermentation stresses. This phenotype allows growth in crude hydrolysates, where inhibitory substances such as aldehydes are present and might inhibit growth and xylitol production, consequently (Albuquerque et al., 2014).

Finally, we carried out an electro-fermentation to evaluate the effect of using such system on xylose conversion. To address the limitations of traditional fermentations, electro-fermentation has been studied as an emerging strategy that influences microbial metabolism, product formation and even their recovery. This bioprocess makes use of electrodes that function as electron acceptors, which interact with microorganisms and allow the formation and exchange of electrons. Electrochemical processes improve pH control, redox balance, optimizes carbon source, increases ATP synthesis, and improve the yield of cell biomass, among other advantages. Such methodology have already been used with residual raw material from the food industry for other applications (Chandrasekhar et al., 2021; Rago et al., 2019; Schievano et al., 2016).

Together with an initial OD_600_ of 2.5 and hydrolysate supplementation, 1 volt of electrical current was applied for the fermentation of sugarcane hydrolysate. Such electric tension was chosen due to previous internal results obtained with the same system for improved fermentation (results not published). The system available to test such condition is shown in **Figure 2A**. At the end of this process, FMYX was able to produce 6.72 ± 0.40 g/L of xylitol, while the control without electric current resulted in 4.34 ± 0.31 g/L of the reduced sugar (**Figure 2B** and **2C**, respectively): representing an improvement of 54.8% in xylitol production and a productivity of 0.046 g/L for the last. Nevertheless, a low yield of xylitol was obtained, possibly due to the aeration conditions of such process. The use of such bioelectric cell did not allow for the use of Erlenmeyers – as it can be observed in **Figure 2A** -, which would create better oxygenation conditions. The use of a Becker was mandatory given the material and structure available for this assay, and better aeration could not be provided for testing. Still, it was possible to observe enhanced xylitol production when the electric current was added to the fermentation process. Supplying electrical impulses during fermentation allows better availability of cofactors that improve yeast metabolism (Moscoviz et al., 2016), which could explain the increase in xylitol production.

For all the fermentation scenarios previously described a notable decrease in xylose concentration for FMYX occurred after 50 h of fermentation, accelerating towards the end of the process. Total depletion of glucose was observed in the first 20 h of cultivation. Once the depletion of glucose occurred, the ethanol produced was assimilated by the yeast for the regeneration of cofactors, since its concentration decreases at the end of the fermentation for the industrial strain - a similar effect was reported by Baptista et al., (2018). The highest titer of xylitol produced was 19.24 ± 0.68 g/L, representing a productivity of 0.132 g/L.h, achieved by FMYX and combining a higher load of initial cell density and supplementing the hydrolysate with peptone and yeast extract.

Better xylitol productivity results have been reported when using other biomass hydrolysates and recombinant *S. cerevisiae*, but with different fermentation strategies. (Baptista et al., 2018) was able to produce 148.5 g/L, while Kogje and Ghosalkar, (2017) efforts resulted in 47 g/L xylitol production. Nevertheless, both strategies used a fed-batch approach for xylose metabolization while this report focuses on batch fermentation. The last author also used detoxified hydrolysate, a requirement that would imply further manipulation of the sugarcane straw and elevate process costs. Again, non-fed cultivation of non-detoxified biomass is a proper alternative for xylitol production in an industry that already applies such system – in special the second-generation sugarcane bioethanol production –, and that is the reason it was applied here.

A challenge that still needs to be optimized in the bioproduction of xylitol is the purification and recovery of the product at the end of the process. An expensive step that compromises the large-scale application, due to impurities, by-products and compounds generated in the hydrolysate during fermentation. To achieve a high degree of purity of xylitol, the combination of several traditional methodologies is required, including the use of activated carbon, ion exchange resins, chromatography, among others. However, the percentages of recovery reveal a partial loss of the sweetener, for which the optimization of the steps prior to production could facilitate its recovery and purification (Queiroz et al., 2022).

## 4. CONCLUSION

The application of our strategies in this study are important from an economic perspective because it indicates the possibility of optimizing the fermentation process for the efficient use of renewable feedstock and xylitol production using recombinant *S. cerevisiae* as a biocatalyst. Our work provides evidence to establish a less expensive alternative process for xylitol production, easily applicable at an industrial level, that can take advantage of the already established E2G infrastructure. As our results highlighted, the strategy of increasing the initial cell density and medium supplementation showed a relevant increase in the final titer of xylitol. Also, it is evidenced that the use of an electro-fermentation improves xylitol production, and better aeration conditions are suggested for better results. This is one of the few reports on xylose fermentation using sugarcane straw hydrolysate focused on the production of xylitol, and not exclusively for ethanol, using *S. cerevisiae*, and also the first to apply a bioelectric system for this purpose.

## ACKNOWLEDGEMENT

The authors are grateful for the funding provided by the Brazilian research agencies CAPES (Coordenação de Aperfeiçoamento de Pessoal de Nível Superior), and FAPESP (Fundação de Amparo à Pesquisa do Estado de São Paulo, 2021/02421-0).

